# Colibactin-producing *E. coli* promote carcinogenesis of gastroesophageal adenocarcinoma and simultaneously induce autophagy and differentiation

**DOI:** 10.1101/2025.11.10.687579

**Authors:** Andrea Proaño-Vasco, David Obwegs, Kerstin Bruder, Céline Ritzkowski, Sophia Eichner, Lauren Houle, Yiwei Sun, Ceylan Tanes, Lioba Klaas, Linus R. Schömig, Donja Sina Mohammad-Shahi, Kyle Bittinger, Kenneth K. Wang, Anil K. Rustgi, Gary W. Falk, Robert Thimme, Timothy C. Wang, Harris H. Wang, Bärbel Stecher, Ulrich Dobrindt, Julian A. Abrams, Sagar, Michael Quante

## Abstract

**Background & Aims:** Gastroesophageal adenocarcinoma (GEAC) is a malignancy of the gastroesophageal junction (GEJ) and is associated with reflux of gastroduodenal contents and Barrett’s Esophagus (BE). A shift towards gram-negative bacteria in the microbiota of the GEJ additionally promotes inflammation and likely carcinogenesis. *Enterobacteriaceae* are enriched in advanced stages of GEAC development, and members of this family can produce colibactin, a genotoxin implicated in DNA damage and tumor progression. We aimed to validate these observations and investigate the effect of *E. coli* with or without colibactin production on GEAC-carcinogenesis.

**Methods:** Bacteria were profiled in human biopsies with imaging and 16S rRNA gene sequencing. Organoids of our L2-IL1B mouse model of GEAC were exposed to *E. coli* with colibactin (CoPEC) and without colibactin production (noCoPEC) via organoid microinjection. The phenotypic and transcriptomic changes in the organoids after the coculture with *E. coli* were evaluated via histology and single-cell RNA sequencing.

**Results:** In human specimens, we observed an infiltration of bacteria in GEJ-tissue upon tumor formation and detected *Enterobacteriaceae* in one third of BE-patients. CoPEC-injected organoids exhibited high rates of proliferation and DNA damage, and an upregulation of cancer-associated genes and pathways. Furthermore, genes and pathways associated with immune activation, defense mechanisms, metabolic reprogramming, autophagy and differentiation were upregulated in CoPEC-injected organoids.

**Conclusion:** In addition to the enrichment of *Enterobacteriaceae* in the GEJ-tissue of patients at late stages of GEAC, we show that the exposure of colibactin-producing *E. coli* to murine BE-organoids promotes genetic instability and proliferation, and the activation of cancer-associated pathways, while also activating autophagy and enhancing intercellular homeostasis. This indicates that colibactin-producing *E. coli* have a dual effect on early stages of GEAC-carcinogenesis.

**GRAPHICAL ABSTRACT:** 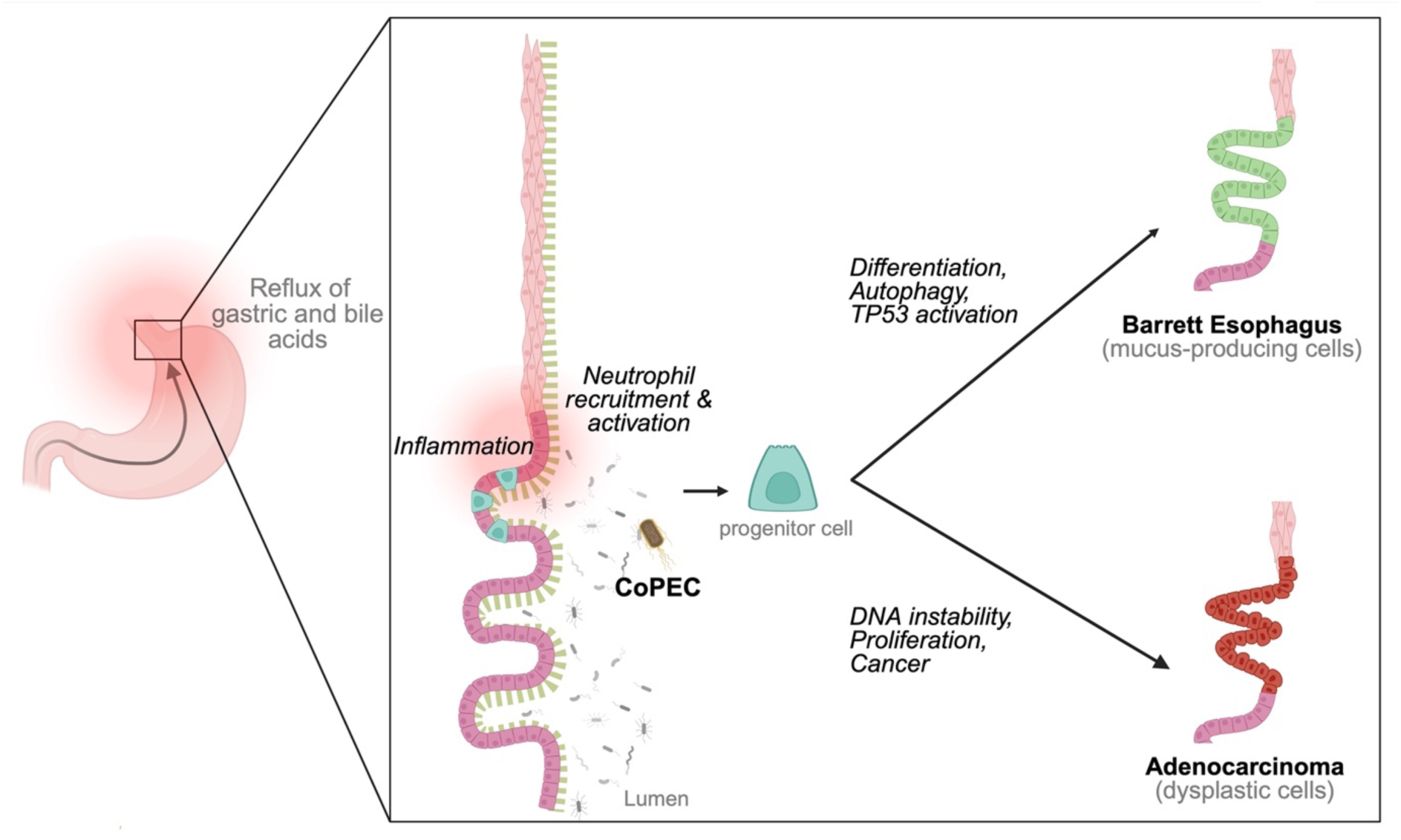

## INTRODUCTION

Gastroesophageal adenocarcinoma (GEAC), comprising esophageal and gastroesophageal junction adenocarcinoma, is prevalent in high-income Western countries. It predominantly affects men and is associated with poor survival rates, particularly among older patients^1^. A western-style diet, characterized by a high intake of saturated fats and refined sugars, is linked to obesity and gastroesophageal reflux disease (GERD), the strongest risk factor for GEAC^2^. This results in a chronic exposure of the epithelium at the gastroesophageal junction (GEJ) to gastric and bile acids. Such exposure compromises the integrity of the protective mucus barrier, induces shifts in the local microbial community, and increases the direct interaction of epithelial cells with pro-inflammatory and genotoxic compounds. Although genetic, environmental, and lifestyle risk factors have been well characterized, the contribution of the microbiota to GEAC pathogenesis remains poorly understood. Emerging evidence suggests that shifts in the microbiota at the GEJ influence carcinogenesis through metabolic and inflammatory pathways^3, 4^. However, the links between esophageal pathologies and the microbiota remain largely correlative, and the identification of specific microbes driving disease progression is still in its early stages^3^. Further research in this area could support the development of novel biomarkers, chemoprevention through targeted antimicrobial therapies, and predictive tools for disease outcome.

The gastrointestinal tract hosts a complex and diverse microbial community, with the colon being the most densely populated organ. Although less abundant, the esophagus also hosts a local microbiota, which was first characterized in 2004 and revealed a microbial community distinct from that of the oral cavity^5^. Subsequent studies have confirmed these findings^3, 6^, and demonstrated the predominance of gram-positive bacteria in the healthy esophageal mucosa^7, 8^. Existing studies suggest that microbial composition undergoes substantial alterations during GEAC-carcinogenesis^3, 4^ and support the notion that tumor formation is facilitated by dysbiosis of the local microbiota. Indeed, the esophageal microbiota of patients with reflux esophagitis or Barrett’s esophagus (BE), a metaplastic lesion associated with GEAC, exhibited an increased abundance of gram-negative bacteria (GNB)^7, 9, 10^, which are known to promote inflammation via the lipopolysaccharides (LPS) on their outer membrane^11^. Within GNB, the bacterial family *Enterobacteriaceae* has been found to be increased in advanced stages of GEAC^4^. Notably, some members of this family produce colibactin, a genotoxin known to cause DNA double-strand breaks leading to cell cycle arrest, chromosomal instability, senescence *in vitro*, and tumorigenesis *in vivo*^12^.

Given the shift from gram-positive to gram-negative bacteria at the GEJ with gastroesophageal reflux and BE, upon disease development, the increased abundance of *Enterobacteriaceae* at late stages of GEAC development^4^, and the fact that colibactin-producing *Enterobacteriaceae* have been implicated in colon cancer^13^, we sought to investigate the effects of *E. coli* (noCoPEC) and colibactin-producing *E. coli* (CoPEC) on GEAC-carcinogenesis using specific *E. coli* strains as model organisms. We isolated organoids from the mouse model of BE and GEAC (L2-IL1B)^14^ and exposed them to CoPEC, noCoPEC, or food colorant (FC) via microinjection. The resulting phenotypic alterations and the transcriptomic profile of the organoids were evaluated via histology and single-cell RNA sequencing (scRNA-seq), respectively.

## RESULTS

### High relative abundance of gram-negative bacteria in Barrett’s esophagus

Previous studies of esophageal microbiota composition could not determine whether bacteria were biologically relevant or represented superficial “bystanders” in the esophageal lumen. To assess whether bacteria could play a role in esophageal carcinogenesis, we examined BE biopsies for bacterial presence either closely adherent to or within the tissue. We collected biopsies during upper endoscopy in 14 BE patients. In order to preserve the mucus layer, we fixed the biopsies in methacarn prior to paraffin embedding. We visualized the bacteria and mucus layer using fluorescence *in situ* hybridization and WGA staining, respectively, followed by microscopy. Interestingly, we found bacteria within the stroma of BE-tissue but only minimal bacteria within the surface mucus layer (Figure 1A-B). Importantly, we found an increase in bacterial density in biopsies from patients with high-grade dysplasia or intramucosal adenocarcinoma as compared to no dysplasia, indefinite for dysplasia, or low-grade dysplasia (Figure 1C).

**Figure 1:**
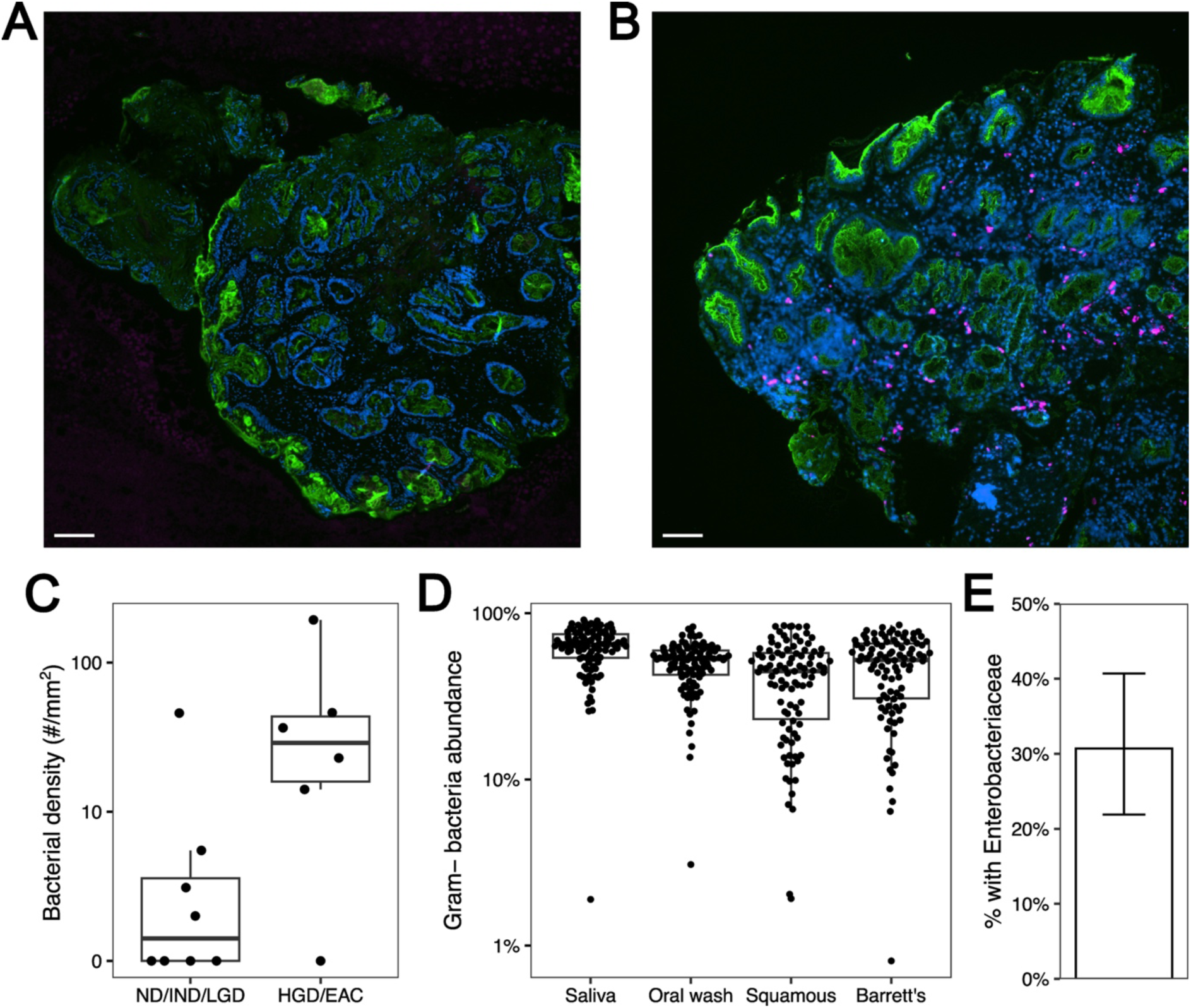
Bacterial presence and characterization in Barrett’s esophagus Fluorescence *in situ* hybridization (FISH) images of biopsies from (A) BE-patient without dysplasia and (B) BE-patient with intramucosal adenocarcinoma. Stains: bacteria (pink, Eubacteria), mucus layer (green, WGA), nuclei (blue, DAPI). White bars represent 100 µm. (C) Increased bacterial density in BE with HGD/ EAC (n=6) vs. ND/IND/LGD (n=8) (p=0.056). (D) High relative abundance of gram-negative bacteria in BE subjects (n=101) across saliva, oral wash, and squamous and BE tissue brushings. (E) *Enterobacteriaceae* were detected in BE tissue in 30.7% of subjects (n=101).

We also re-analyzed 16S rRNA sequencing data from a multi-center cross-sectional study of patients from the BETRNET consortium without and with Barrett’s esophagus with biospecimens collected at the time of upper endoscopy^15^. While not statistically distinct from non-BE controls, we found high relative abundances of gram-negative bacteria across saliva, oral wash, and squamous and BE tissue brushings (Figure 1D). Furthermore, we detected *Enterobacteriaceae* in BE tissue in 30.7% of patients (Figure 1E).

### Microinjection of organoids with bacteria to model microbe-host interactions at the gastroesophageal junction

We successfully established a microbe-organoid coculture system based on the protocol developed by Puschhof et al., 2021^16^, and adapted it for application in BE- and GEAC-research. Using this system, we examined the direct interactions between CoPEC and epithelial cells derived from dyplastic BE-tissue in a L2-IL1B mouse (mouse model of BE and GEAC)^14^, resembling LGD in humans. Briefly, two independent rounds of microinjection were performed on organoids from the same mouse line. In both rounds, the *E. coli* strain producing colibactin (M1/5 *rpsL*K42R-*gfp*, CoPEC), the control *E. coli* strain (M1/5 *rpsL*K42R-*gfp* Δ*clbR*, noCoPEC), and food colorant (FC) were microinjected into the lumen of the organoids to model direct microbe-host interactions. Organoids from the first microinjection round (passage 7) were dissociated into single cells and sorted into 384-well plates for scRNA-seq to assess transcriptomic changes at single-cell resolution. Organoids from the second round (passage 10) were fixed, embedded in paraffin, and subjected to histological analysis to evaluate phenotypic alterations (Figure 2A). This integrated approach allowed us to characterize both morphological and molecular responses of epithelial cells from organoids derived from dyplastic BE-tissue to CoPEC-exposure, providing insight into microbial contributions to neoplastic transformation.

**Figure 2:**
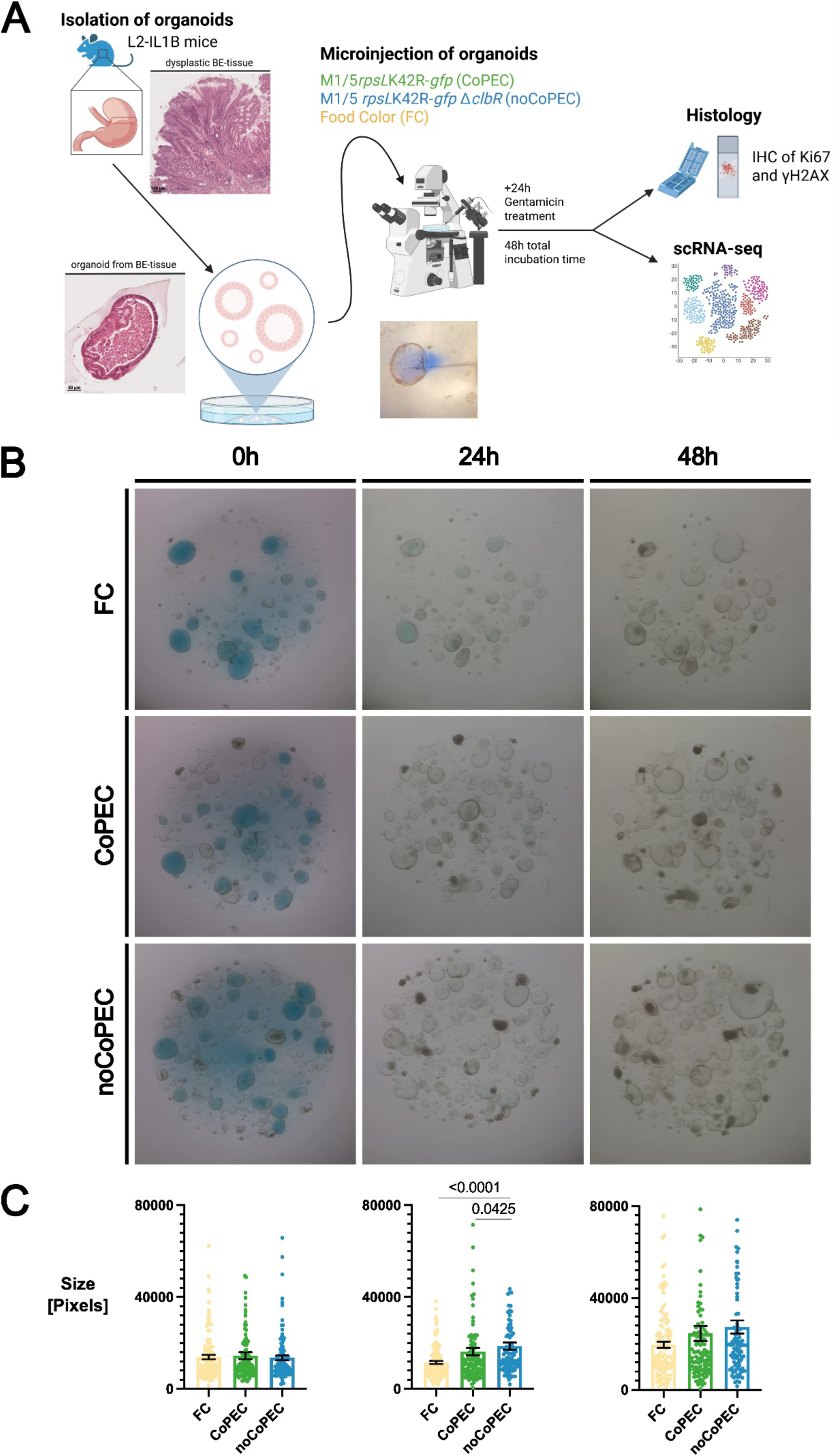
Experimental setup for microinjection of organoids with bacteria and downstream analyses, as well as macroscopic evaluation of the organoids (A) Graphical representation of the microinjection of organoids with bacteria and food colorant: Organoids were isolated from 9-month-old L2-IL1B mice and kept in culture during multiple passages (7-10) until reaching organoid size and confluency needed for the microinjection. Histological images depict the dysplastic BE-tissue of the mouse the organoids were isolated from, and an exemplary dysplastic organoid. For microinjection, organoids were seeded in a 4-well plate and were microinjected with either a blue-stained solution containing colibactin-producing E. coli (CoPEC), a blue-stained solution containing colibactin-non-producing E. coli (noCoPEC), or the bacteria-free blue-stained solution 4 days after the last passage. After microinjection, pictures of the wells were taken, and the growth media was exchanged for gentamicin-containing (10 µg/mL) growth media. Phenotype of the organoids was evaluated via histological analysis (IHC of Ki67 and γH2AX). Genetic alterations in the organoids were examined via scRNA-seq. (B) Representative macroscopic images from the microinjection round for scRNA-seq showing organoids injected with FC, CoPEC and noCoPEC at 0h, 24±4h, and 48h±4h. Images of every well were taken right after microinjection and every 24±4h. (C) Evaluation of the macroscopic images. The success rate of injection was assessed by dividing the area of injected organoids by the total area of all organoids in the well. Average success rate of injection from the microinjection round for scRNA-seq at passage 7 and for histology at passage 10: FC (49%), CoPEC (59%), noCoPEC (43.5%). n_FC_= 118 injected organoids, 16 wells, 2 tech. replicates; n_CoPEC_= 107 injected organoids, 15 wells, 2 tech. replicates, n_noCoPEC_= 105 injected organoids, 16 wells, 2 tech. replicates.

### Exposure to *E. coli* enlarges organoid size and increases proliferation rate and DNA damage

Macroscopic evaluation revealed that 24h after exposure, organoids injected with bacteria were larger than those injected with food colorant (Figure 2B-C). Increased proliferation and genomic instability are recognized as hallmarks of cancer^17^. These features were assessed histologically by quantifying Ki67-positive and γH2AX-positive cells, respectively. Although both bacterial strains affected the organoids, CoPEC-organoids exhibited a higher proliferation rate and significantly higher DNA damage compared to FC-organoids (Figure 3A-B). Furthermore, analysis of the cell cycle state at transcriptomic level showed CoPEC-injected organoids having the highest proportion of cells in the G2M-phase (Figure 3C). In combination, these results suggest that in CoPEC-injected organoids there were more cells preparing or undergoing cell division.

**Figure 3:**
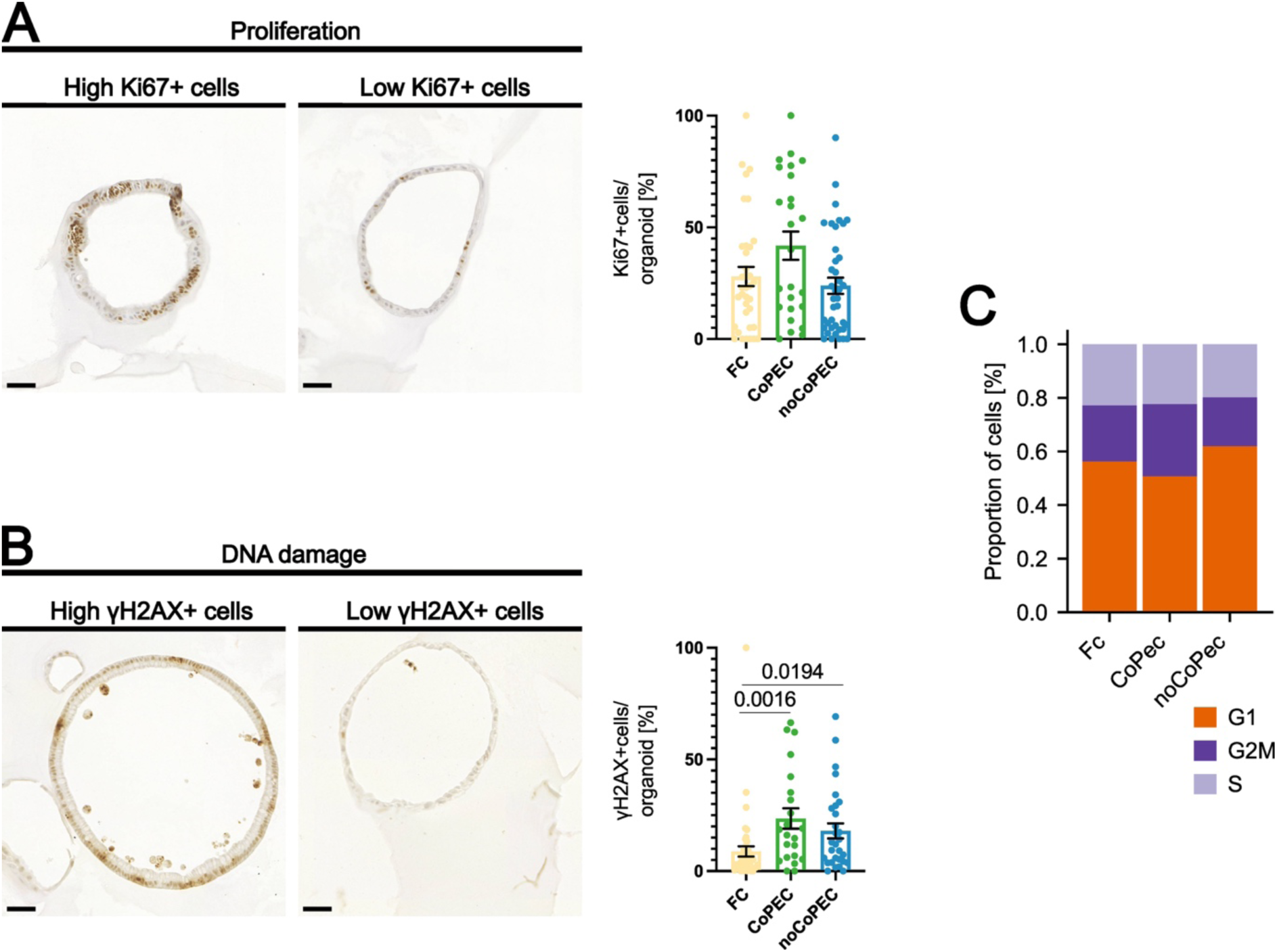
CoPEC-injected organoids showed increased proliferation and DNA damage when compared to noCoPEC- and FC-injected organoids Representative images and evaluation of proliferation rate and DNA damage by immunohistochemistry (IHC) of (A) Ki67+ cells and (B) γH2AX+ cells respectively. Black bars represent 50 µm. (C) Stacked barplot showing the proportion of cells assigned to each phase of the cell cycle as determined by scoring the single cells using published gene sets^103^.

### Bacterial injection into murine BE organoids induces transcriptional signatures associated with immune activation and carcinogenesis

To investigate the transcriptional profiles of organoids injected with CoPEC, noCoPEC and FC, organoids were dissociated into single-cell suspensions and subjected to scRNA-seq (Figure 4). Initial analysis did not reveal distinct clustering of cells by treatment group (Figure 5A). Consequently, single-cell transcriptomes were aggregated by technical replicates into pseudobulks^18^. Principal component analysis (PCA) of the pseudobulked datasets revealed that organoids injected with bacteria (CoPEC and noCoPEC) clustered separately from those injected with food colorant (Figure 5B). Unsupervised differential gene expression (DGE) analysis demonstrated a significant upregulation of genes associated with host-microbe interactions and antimicrobial defense (*Duox2*^19^, *Duoxa2*^20^, *Duoxa1*^21^, *Reg3g*^22^, *Pigr*^23, 24^, *Pglyrp1*^25^, *Lcn2*^26^) in CoPEC- and noCoPEC-injected organoids. Genes involved in inflammatory responses (*Tnfaip2*^27, 28^, *Zc3h12a*^29, 30^, *Pglyrp1*^25^, *Klf6*^31^, *Nfkbia*^32^, *Vnn1*^33^), neutrophil recruitment (*Cxcl1*, *Cxcl3*) and activation (*Cxcl5*)^34, 35^, mucus layer integrity (*Clca3b*^36^), and cancer (*Tnfaip2*^27, 28^, *Slc39a4*^37, 38^, *Plet1*^39^, *St8sia6*^40, 41^, *Vnn1*^33^) were also upregulated in CoPEC- and noCoPEC-injected organoids. Additionally, *Matr3* was significantly upregulated in CoPEC-organoids. *Matr3* is implicated in the regulation of gene expression, DNA damage response, and cellular processes such as proliferation, differentiation and survival^42, 43^ (Figure 5C-E). Collectively, these results show that bacteria-injected organoids exhibit enhanced expression of genes involved in host defense, inflammatory responses, and pathways linked to carcinogenesis.

**Figure 4:**
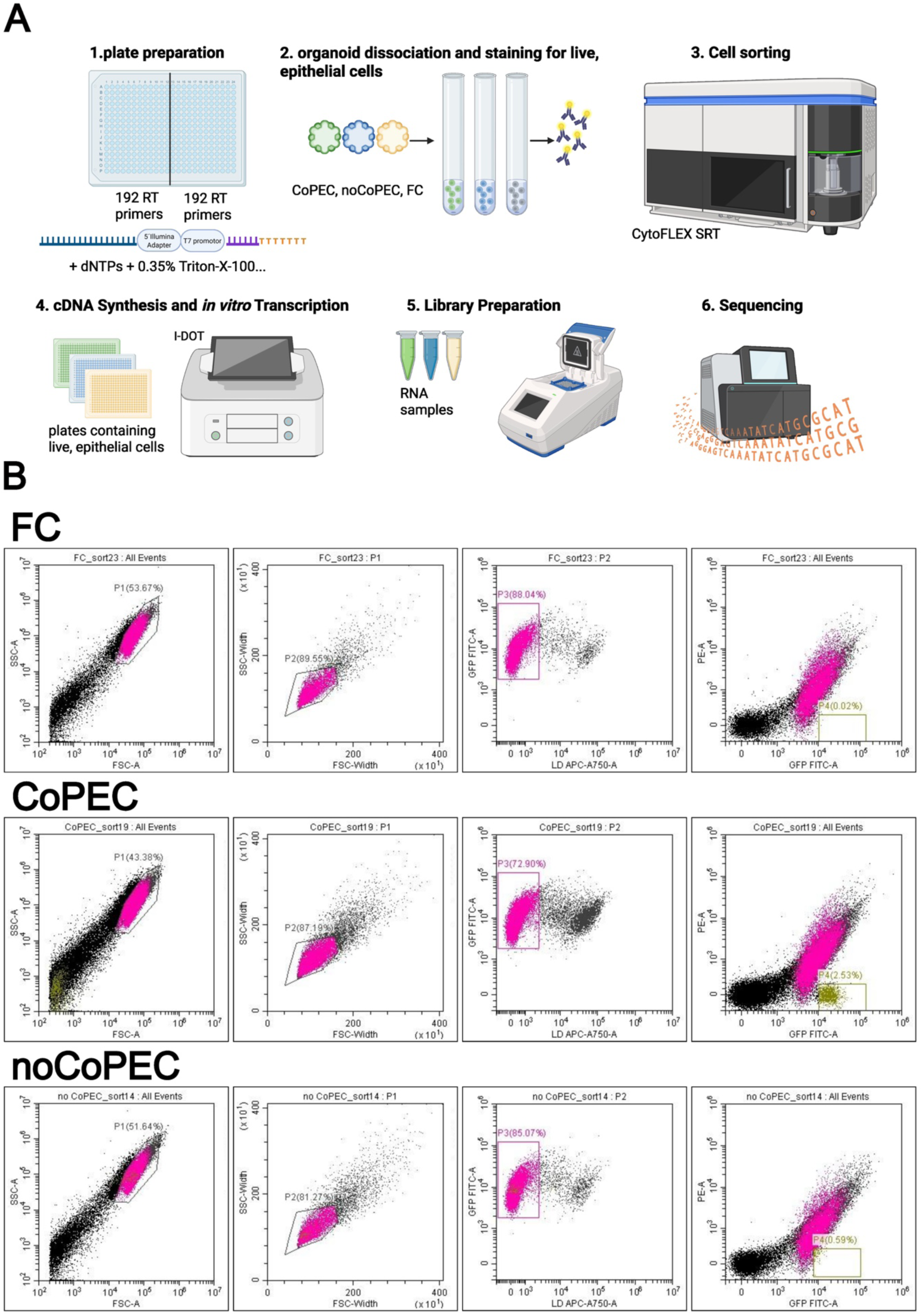
Sorting and scRNA-seq of organoids (A) 1. 384-well PCR plates were prepared by adding mineral oil, lysis buffer and a reverse transcription (RT) primer into each well. Every plate consisted of 2 sets of 192 different RT primers that additionally contained a polyT primer having a 6 bp cell barcode, 6 bp unique molecular identifier (UMIs), a 5’Illumina adapter, and a T7 promoter. 2. Organoids were dissociated into single cells that were then stained with zombie NIR for dead cells. 3. Single, live, epithelial cells were sorted with the CytoFLEX SRT into every well of the previously prepared plate. 4. Reverse transcription, Clean-up for complementary DNA (cDNA), *In vitro* transcription and clean-up for amplified RNA (aRNA). 5. Library preparation, PCR amplification, and PCR clean-up. 6. scRNA-seq. (B) Gating strategy used for defining cells to be sorted in the 384-well plates for scRNA-seq. From the left to the right: first plot was created to differentiate cells based on their size and granularity; second plot was created to identify and exclude cell doublets or aggregates from single cells; third plot was created to exclude cells that were permeant to Zombie NIR^TM^ (APC-A750) and were therefore dead; forth plot was created to check if GFP-positive bacteria were detectable as was the case in CoPEC only.

**Figure 5:**
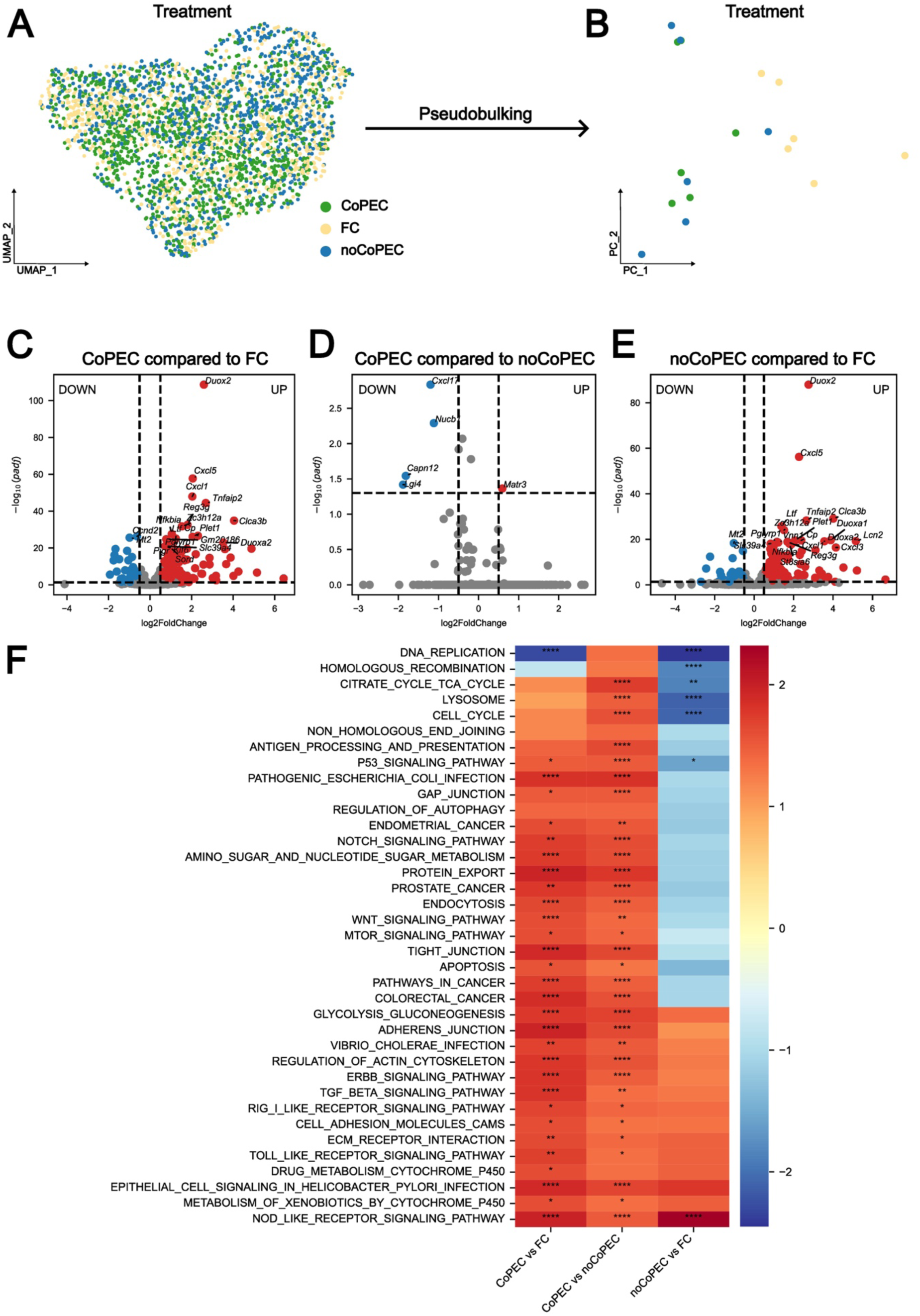
The signature of gene expression and pathway regulation in CoPEC-injected organoids differed significantly from FC-injected and noCoPEC-injected organoids (A) UMAP representation of scRNA-seq data colored by the treatment of the organoids. Every dot represents a single cell. (B) PCA plot showing pseudobulked scRNA-seq data. Here, every dot represents one technical replicate (n=6) colored by treatment. (C-E) Volcano plots resulting from Differential Gene Expression (DGE) Analysis of pseudobulked scRNA-seq data showing overexpression (red dots) and underexpression (blue dots) of genes. Horizontal dashed lines show the adjusted p-value cutoff of 0.05 whereas vertical dashed lines show the log2 fold change threshold of 0.5. (F) Unsupervised gene set enrichment (GSE) Analysis depicting KEGG-pathways on the y-axis and group comparisons on the x-axis of the heatmap. The normalized enrichment score indicates pathway activation if positive, or pathway suppression if negative, and it is depicted as bar on the right. * ≤ 0.05, ** ≤ 0.01, *** ≤ 0.001, **** ≤ 0.0001.

### CoPEC injection led to enhanced activation of metabolic, oncogenic, and cellular defense pathways in organoids

Unsupervised gene set enrichment (GSE) Analysis revealed significant upregulation of the NOD like receptor (NLR) signaling pathway in both CoPEC- and noCoPEC-organoids (Figure 5F). NLRs are intracellular sensors that detect microbial infection and cellular stress^44^. In CoPEC-injected organoids several additional pathways were significantly upregulated. These included metabolism-associated pathways such as amino sugar and nucleotide sugar metabolism, glycolysis, and gluconeogenesis; housekeeping pathways including mTOR signaling^45^, endocytosis and protein export^46^; and pathways involved in tissue homeostasis and cell communication such as ErbB signaling^47^, regulation of actin cytoskeleton^48^, ECM receptor interaction, cell adhesion molecules, adherens junctions, tight junctions, and gap junctions^49^. The TGFβ signaling pathway, which was also upregulated in CoPEC-injected organoids, has diverse, context-dependent outcomes that include inhibition or promotion of proliferation, apoptosis, autophagy and senescence^50^. Moreover, bacteria-driven alterations of epithelial cells including epithelial cell signaling pathways associated with *Helicobacter pylori*, pathogenic *E. coli*, and *Vibrio cholerae* infections, were significantly enriched in CoPEC-injected organoids. Likewise, pathogen recognition pathways such as RIG-I like receptor^51^ and Toll-like receptor signaling^52^, together with cellular defense mechanisms including apoptosis^53^ and metabolism of xenobiotics by cytochrome P450^54^, were significantly upregulated in CoPEC-injected organoids. Notably, Notch- and Wnt-signaling pathways, as well as cancer-associated pathways, were also significantly upregulated in CoPEC-injected organoids. Furthermore, the cell cycle, citrate cycle, lysosomal pathway, and p53 signaling pathway were all significantly upregulated in CoPEC-injected organoids but significantly downregulated in noCoPEC-injected organoids (Figure 5F). Together, these results suggest that CoPEC specifically drives broad transcriptional changes in organoids, activating metabolic, signaling, and cancer-related pathways while enhancing epithelial defense and tissue organization.

### Colibactin-producing *E. coli* induces increased expression of stem cell-, epithelial-, autophagy-, and cancer-related genes

Targeted analysis of scRNA-seq data for selected gene sets related to cell phenotype and plasticity showed a significantly increased expression of genes associated with stem cells and epithelial cells in CoPEC-injected organoids compared to the other treatment groups. Compared to noCoPEC-only, CoPEC-injected organoids showed a significantly increased expression of genes linked to epithelial differentiation and cell plasticity (Figure 6A-B). Consistent with the histological observations (**Error! Reference source not found.**), the targeted analysis of scRNA-seq data for selected functional genes demonstrated a significantly elevated expression of genes associated with proliferation and cell division, as well as DNA damage and repair in CoPEC-injected organoids compared to noCoPEC-injected organoids. The expression of genes associated with autophagy, a process that mediates the degradation of intracellular components^55^, was also significantly enhanced in CoPEC-injected organoids only (Figure 6C-D). Furthermore, CoPEC-injected organoids showed an increased expression of genes that, if overexpressed or dysregulated, are linked to cancer. These included genes associated with metabolic (*Iars2*^56^*, Oxct1*^57^*, Hmgcs1*^58^*, Acsl4*^59^*, Prdx1*^60^*, Idh1*^61^*, Acadm*^62^*, Ldha*^63^*, Slc25a17*^64^*, Aldh3a1*^65^), and signaling (*Kras*^66^*, Gnaq*^67^*, Src*^68^*, Grb2*^69^*, Cd38*^70^*, Erbb3*^71^*, Bmp2*^72^) pathways (Figure 6E-F). Together, these results suggest that the exposure to CoPEC promotes stem-like and differentiated cell phenotypes, enhances cellular defense mechanisms such as autophagy and DNA repair, and activates metabolic and oncologic signaling pathways associated with cancer development.

**Figure 6:**
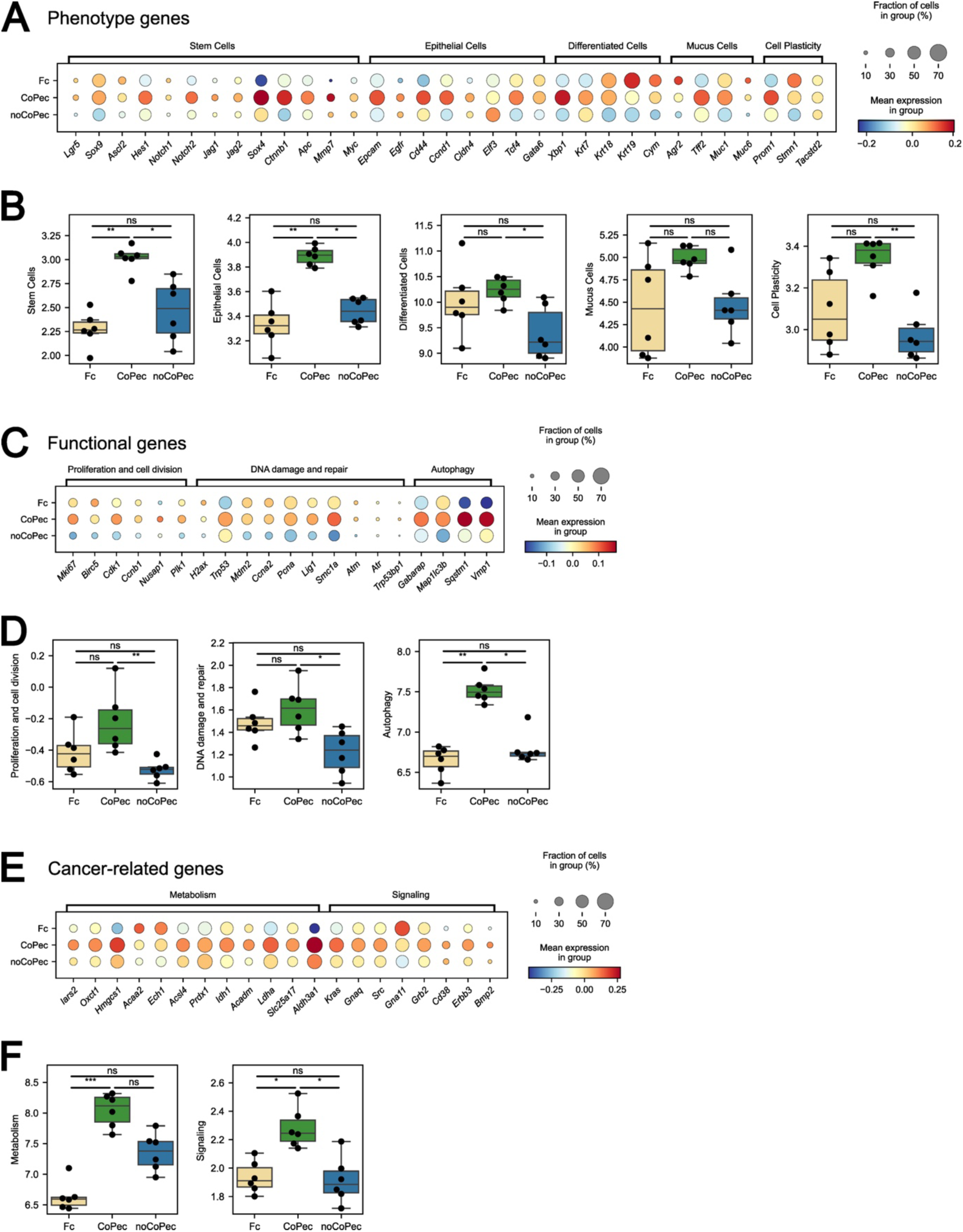
The exposure to colibactin-producing *E. coli* increases the expression of stem cell-, epithelial-, autophagy-, and cancer-related genes (A) Dot plot of selected phenotype genes. (B) Boxplots comparing the expression of gene sets related to cell phenotypes. (C) Dot plot of selected functional genes. (D) Boxplots comparing the expression of gene sets related to cellular functions. (E) Dot plot of selected genes related to metabolic and signaling genes. (F) Boxplots comparing the expression of gene sets involved in metabolic and signaling pathways, that if dysregulated, are associated with cancer. (A, C, E) Dot color represents scaled transcription values and dot size shows proportion of cells expressing the genes. (B, D, F) Each technical replicate is a dot. Score was calculated using a Univariante Linear Model (ULM). Statistical testing was performed using Kruskal-Wallis and Dunńs post hoc test with Holm adjustment for multiple testing. * ≤ 0.05, ** ≤ 0.01, *** ≤ 0.001, **** ≤ 0.0001.

## DISCUSSION

GEAC is a malignancy of the GEJ^1^ and is associated with a western-style diet, obesity, reflux of gastroduodenal contents and Barrett’s Esophagus (BE)^2^. Chronic exposure of the epithelium to gastric and bile acids compromises the integrity of the protective mucus barrier, induces shifts in the microbial community at the GEJ, and increases the direct interaction of epithelial cells with pro-inflammatory^73^ and genotoxic compounds^74^. Although the contribution of microbes to GEAC pathogenesis remains poorly understood, emerging evidence suggests that shifts in the microbial composition at the GEJ influence carcinogenesis through metabolic and inflammatory pathways^3, 4^. Here, we demonstrate the ability of colibactin-producing *E. coli* to alter cellular metabolism, gene expression signatures, and phenotype in a direct interaction with metaplastic epithelial cells.

The increased bacterial density in biopsies from patients with high-grade dysplasia and GEAC supports previous observations indicating a shift towards gram-negative bacteria in the microbiota at the GEJ of patients with reflux esophagitis and BE^7, 9, 10^. *Enterobacteriaceae,* belonging to the gram-negative bacteria with the ability to produce colibactin^12^, were detected in GEJ-tissue of one-third of BE-patients in our patient cohort, as previously observed^4^. Given the implication of colibactin-producing *Enterobacteriaceae* in colon cancer development^13^, we explored the effect of colibactin-producing *E. coli* on GEAC-development by exposing BE metaplastic mouse organoids to CoPEC, noCoPEC, and FC via microinjection.

Indeed, the transcriptomic profile of the organoids exposed to *E. coli* revealed a coordinated response to bacterial infection characterized by the overexpression of genes associated with host-microbe interactions and defense mechanisms against pathogens, inflammation, neutrophil recruitment, and activation. This is consistent with prior observations that exposure of epithelial cells to enterotoxigenic (ETEC), enteropathogenic (EPEC) and enterohemorrhagic (EHEC) *E. coli* induces the expression of genes linked to inflammatory pathways and metabolic reprogramming^75, 76^.

Notably, the expression of autophagy-related genes was also upregulated in CoPEC-infected organoids. Autophagy, a self-degradative process crucial for preserving cellular homeostasis, functions as a key innate immune mechanism by selectively clearing intracellular pathogens and is activated during diverse bacterial and viral infections. Beyond pathogen clearance, autophagy prevents malignant transformation by eliminating damaged organelles, degrading misfolded proteins, and limiting oxidative stress^77^. Conversely, it may also foster carcinogenesis by promoting tumor cell survival and proliferation under metabolic and hypoxic stress^78^. In human CRC cells, CoPEC infection induces autophagy as a protective response aimed at eradicating intracellular bacteria^78^. In line with these observations^78^, our results suggest that autophagy is a key mechanism by which cells respond to infection by colibactin-producing *E. coli*.

In addition, *Matr3*, a gene that encodes for an A/T-rich DNA-binding nuclear scaffold protein^42^, is a target of ATM and is implicated in the early stage of the cellular response to DNA double-strand damage^79^. P53 is activated in response to many stress signals such as the ATM-dependent response to DNA damage^80^. Recently, it was elucidated that colibactin alkylates DNA at A/T-rich sites^81^. Accordingly, the distinct overexpression of *Matr3* and *Tp53* found only in CoPEC-injected organoids points to a host defense mechanism specifically triggered by colibactin.

Analysis of scRNA counts revealed an overexpression of genes linked to stem cells and epithelial cells, as well as to differentiation and cell plasticity in CoPEC-injected organoids. Notch-and Wnt-signaling pathways play a key role in stem cell maintenance^82, 83^. In the L2-IL1B mouse model, activation of the Notch signaling in epithelial cells has been shown to promote disease development, whereas its inhibition mitigated the disease phenotype. Additionally, Iftkhar et al., 2021 reported an increased expression of Wnt target genes in colon organoids infected with colibactin-producing *E. coli*^83^. Therefore, the elevated expression of stem-cell related genes and the upregulation of these pathways in CoPEC-injected organoids observed in this study, suggest an enhanced stem cell activity upon infection with colibactin-producing *E. coli.* Progenitor cells give rise to specialized epithelial lineages, including mucus-producing cells^84^. The resulting mucus layer forms a dynamic protective barrier that represents the first line of innate immune defense^85^. Indeed, it was demonstrated that the adherent mucus layer on epithelial cells mitigates the genotoxic effects of colibactin^86^. Consistently, the mucus-associated genes *Muc1* and *Tff2*^87, 88^ were overexpressed in CoPEC-organoids, suggesting an activated host defense response. The highly organized structure and polarity of the epithelium is maintained by cell junctions that connect adjacent cells, and control substance diffusion and allow selective permeability^89^. Epithelial integrity is one of the innate defense mechanisms the gastrointestinal epithelium^90^ and hence, the observed significant upregulation of pathways related to tissue architecture and intercellular communication in CoPEC-injected organoids, suggests a coordinated collective epithelial response aimed at maintaining integrity under bacterial challenge.

CoPEC-injected organoids exhibited increased proliferation and DNA damage, accompanied by an accelerated cell cycle. In addition to the increased expression of genes linked to cancer if overexpressed or dysregulated, CoPEC-injected organoids showed an upregulation of pathways related to prostate and endometrial cancer and cell proliferation. Estrogen metabolites, particularly catechol estrogens, can be oxidized and interact with DNA, indirectly causing mutations and potentially initiating the development of different cancers^91, 92^. Given the genotoxic activity of catechol estrogens^91, 92^ and colibactin^93^, these findings emphasize the contribution of genotoxic agents to dysplasia development.

While our work provides valuable insights linking colibactin-producing *E. coli* to GEAC-carcinogenesis, some limitations have to be acknowledged. As we used organoids derived from the mouse model of BE and GEAC, particular care must be taken when extrapolating the results to humans. Specifically, due to the anatomical differences between mice and humans, and because the environmental conditions are substantially different. Given that the organoid lumen is the site naturally accessible to the environment and microbes, the microinjection of bacteria into the organoids nicely mimics the physiological setting. However, the microinjection method is complex and cannot be performed on a high-throughput basis. Considering that this work aimed to uncover the mechanistic causality between colibactin-producing *E. coli* and GEAC-carcinogenesis, a single biological replicate and multiple technical replicates were used. In our human samples, 16S rRNA gene sequencing was performed to characterize the microbiota, and thus we were unable to specifically assess for the presence of pks+ *Enterobacteriaceae*. Therefore, further studies using human samples and including more biological replicates are needed to validate our observations.

In summary, gram-negative bacteria and especially colibactin-producing *E. coli* can disrupt epithelial homeostasis at the GEJ by imposing a dual pressure on early progenitor cells. On the one hand, bacterial toxins, metabolites, and inflammatory signalling activate autophagy, DNA damage responses, and lineage reprogramming, thereby driving differentiation toward mucus-secreting lineages that reinforce barrier function and represent a protective metaplastic adaptation. On the other hand, persistent colibactin-induced DNA damage, incomplete repair, and proliferative signalling override protective checkpoints, enabling progenitor cells to expand under stress and accumulate mutations. We suspect that this microbial influence thus biases progenitor cell fate decisions between a protective, metaplastic trajectory and a dysplastic, carcinogenic trajectory, establishing bacteria as critical modulators of epithelial plasticity and early carcinogenesis in Barrett’s esophagus and gastroesophageal adenocarcinoma.

## CONCLUSION

Studies investigating microbial dynamics during GEAC-carcinogenesis have shown shifts in the esophageal microbiota^3 4^, supporting the hypothesis of dysbiosis contributing to tumor development. In this study, we successfully reproduced the microbe-epithelium coculture protocol established by Puschhof et al., 2021^16^ and extended its application to BE and GEAC research. Using this model, we examined direct interactions between colibactin-producing *E. coli* and epithelial cells derived from the mouse model of BE and GEAC. Our findings reinforce the hypothesis of colibactin-producing *E. coli* actively contributing to carcinogenesis by promoting proliferation, DNA damage, and the activation of metabolic and signaling pathways associated with cancer, while simultaneously activating defense mechanisms against pathogens, inducing autophagy and enhancing tissue homeostasis. Furthermore, this work supports the emerging idea of GEAC developing as a result of an inflammation-driven expansion of progenitor cells at the gastroesophageal junction, which undergo a fate decision that leads to either the development of Barrett’s Esophagus or to GEAC^82, 94^. Despite its limitations, this model provides a valuable platform for investigating how luminal exposure of bacteria impacts epithelial tissues and contributes to cancer development through microbial metabolites.

## MATERIALS AND METHODS

### Bacterial density assessment of human biopsies

Patients with and without Barrett’s esophagus (BE) scheduled to undergo upper endoscopy at Columbia University Irving Medical Center were prospectively enrolled. In BE patients, study biopsies were taken from the mid-Barrett’s segment or from any visible neoplastic lesions (if present). As part of standard clinical care, Seattle protocol biopsies were taken from the BE segment, and visible lesions were removed by mucosal resection. In non-BE controls, biopsies were taken from the gastric cardia within 1 cm of the squamo-columnar junction. BE subjects were categorized based on the highest degree of neoplasia found on any of the biopsies from that exam. For purposes of analysis, BE subjects were categorized as no dysplasia/indefinite for dysplasia/low-grade dysplasia (ND/IND/LGD) or high-grade dysplasia/adenocarcinoma (HGD/EAC). This study was approved by the Institutional Review Board at Columbia University (AAAQ8763).

Biopsies were fixed in Methacarn prior to paraffin embedding to preserve the mucus layer. Fluorescence in situ hybridization was performed using an Eubacteria-specific DNA probe (EUB388-Cy5 probe, Sequence: 5’–/5Cy5/GCW GCC WCC CGT AGG WGT – 3’, Integrated DNA Technologies) to identify bacteria. Biopsies were subsequently stained with DAPI and wheat germ agglutinin (WGA) to identify nuclei and the mucus layer, respectively. Images were obtained using confocal microscopy and analyzed with CellProfiler and ImageJ. Bacterial density was computed by normalizing bacterial counts to tissue area. Bacteria were identified as Cy5+ objects using Otsu thresholding and declumping. Tissue area was determined by Otsu thresholding of the sum of the DAPI and WGA signal, followed by filling of holes and identification of the largest contiguous area. For each slide, a gross total count of bacterial objects was quantified. Each slide’s bacterial object count was then normalized to its respective tissue area, thus resulting in bacterial density represented as the number of bacteria per mm^2^.

### Profiling of gram-negative bacteria and *Enterobacteriaceae* in human samples

16S rRNA gene sequencing data were re-analyzed from a prior study^15^ .In brief, patients with and without BE who underwent upper endoscopy for clinical indications at Columbia University Irving Medical Center, Mayo Clinic-Rochester, and the University of Pennsylvania were prospectively enrolled from February 2018 through September 2021. Patients were >18 years old, and for BE patients, had histologically confirmed BE ≥2 cm in length and took proton pump inhibitors (PPIs) at least daily for 3 months prior to enrollment. Patients without BE were enrolled stratified 1:1 based on current PPI use. The study was approved by the Institutional Review Boards at Columbia University (AAAQ8763), Mayo Clinic-Rochester (17-000574), and the University of Pennsylvania (826888).

Microbiome profiling was performed of saliva, oral rinse, esophageal squamous brushings, and BE (or cardia from controls) brushings, with all specimens collected just prior to or during the endoscopy. This was done by 16S rRNA gene sequencing of the V1-V2 hypervariable regions. DNA was extracted from samples and libraries annealing to the V1-V2 region of the 16S rRNA gene were generated to be sequenced on Illumina MiSeq. Sequence data were processed using QIIME2 version 2019.7. ^95^. Data files from QIIME were analyzed in the R environment for statistical computing. Global differences in community composition were visualized using Principal Coordinates Analysis. Community-level differences between sample groups were assessed using the PERMANOVA test ^96^. The relative abundances of the ASVs that were assigned to the same taxon were summed. The effect of Barrett’s status and PPI use on log-transformed taxon abundances was assessed using linear models. Only the taxa with at least 0.5% mean relative abundance in at least one sample type were tested. When multiple tests were conducted, we adjusted for false discovery rate (FDR) using Benjamini-Hochberg method.

### L2-IL1B mice for organoid isolation

Organoids were isolated from L2-IL1B mice, the transgenic mouse model of Barrett esophagus (BE) and Gastroesophageal adenocarcinoma (GEAC)^14^. The animal experimental work was conducted with the approval of the Regional Council of Baden-Württemberg under the animal experimental permit G-20/175. Mice were kept under specific-pathogen-free (SPF) conditions in the animal facility of the Center for Experimental Models and Transgenic Services (CEMT) at the Universitätsklinikum Freiburg. Mice were housed in groups of up to five animals in individual ventilated cages and were maintained in 12-hour day-to-night cycles. Mice were fed a standard chow diet (KLIBA, NAFAG, 3807) and water *ad libitum* until the dissection date.

### Isolation of organoids from L2-IL1B mice

L2-IL1B-mice were euthanized by an isoflurane (798-932, CP-pharma) overdose with subsequent cervical dislocation at approximately 9 months of age. Upon dissection, the stomach and esophagus were removed together. The stomach and esophagus were opened from distal to proximal site using scissors. After washing with DPBS (14190-144, Gibco^TM^ by Thermo Fisher Scientific), 1/3 of the cardia (tissue of the gastroesophageal junction) was removed, transferred to a 1.5 µL tube containing 200-300 µL Accutase (A1110501, Gibco^TM^ by Thermo Fisher Scientific) and minced into small pieces. After 15 min incubation on a shaker at RT, tissue was transferred to a 15 mL tube containing 10 mL ice-cold DPBS+0.5M EDTA (AM9260G, Invitrogen by Thermo Fisher Scintific) + 0.5M EGTA (3050.2, Roth) dissolved in DPBS. After 45 min incubation on ice and constant shaking, tubes were centrifuged at 4 °C and 300 g for 5 min.

Further processing of the tissue was performed in sterile conditions under a laminar flow hood. Supernatant was removed, and the pellet was resuspended thoroughly with 10 mL ice-cold PBS+10%FBS to break down the tissue. Tissue suspension was then passed through a 70 µm cell strainer (352350, FALCON®) into a 40 mL collection tube. For a higher extraction yield, the remaining tissue on the strainer was gently pressed against it with the plunger of a syringe. The strainer was washed twice with wash buffer on top of the collection tube. After centrifugation at 4 °C and 300 g for 5 min, removal of the supernatant, resuspension of the pellet in Matrigel® Matrix (MG, 356231, Corning), 30 µL-domes of MG-cell-suspension were distributed in single wells of a 24-well plate (833922, SARSTEDT). When the MG-cell-domes solidified (after 5-10 min), growth medium was added. Organoids were kept in an incubator (37 °C and 5 % CO_2_). Growth medium was changed every 2-3 days and depending on the growth rate, organoids were passaged every 5-7 days.

Maintenance of the organoids (change of growth medium and passaging) was performed in sterile conditions under a laminar flow hood. When passaging the organoids, 100-300 µL DMEM+++ (Advanced DMEM F12 with 1% P/S, 1% HEPES, 1% GlutaMAX^TM^) were added to each well. The content of every well (MG-organoid-domes, organoid media and DMEM+++) was mixed thoroughly and transferred into a 15 mL collection tube (max. 4 wells per tube). Organoids were fragmented by pipetting up and down (5-10x) the content of the tube with a narrow-end Pasteur capillary glass pipette (self-made by melting the tip of the glass pipette with fire, 10230, WU Mainz®). After centrifuging the tubes at RT and 350 g for 5 min, the supernatant was removed, pellets were resuspended in MG, and 30 µL-domes of MG-organoid fragment-suspension were distributed in new wells of a 24-well-plate. The amount of growth medium added to the organoids was dependent on the number of MG domes in every well of the plate and varied between 300-500 µL.

### Preparation of growth medium for organoids based on WRN- and Wnt3A-Medium

Growth medium for organoid culture consisted of self-made 75% L-WRN- and 25% Wnt3a-media with additional growth factors. Following reagents were purchased from Thermo Fisher Scientific: Advanced DMEM F12 (12634010), Value Heat Inactivated FBS (A5256801), Penicillin Streptomycin (P/S, 15140122), HEPES (15630080), GlutaMAX^TM^ (35050061), G418 (10131035), Hygromycin B (10687010), 0.25% Trypsin-EDTA (25200056),B27® (17504001), N2 (17502001),

For WRN (Wnt3A, R-spondin 3 and Noggin) conditioned medium, L-WRN producing cells (ATCC® CRL3276™) were thawed in a water bath at 37 °C, transferred into a 15 mL collection tube previously filled with prewarmed selection medium composed of DMEM-Dil. (Advanced DMEM F12 with 10% FBS, 1% P/S, 1% HEPES, 1% GlutaMAX^TM^), 0.5 mg/mL G418 and 0.5 mg/mL Hygromycin B, and centrifuged at room temperature (RT) and 200 g for 5 min. After discarding the supernatant, the cell pellet was resuspended in 10 mL fresh selection medium, transferred into a prewarmed petri dish (83.3902, SARSTEDT) and maintained in an incubator at 37°C. Medium was exchanged every 2-3 days. When confluent, cells were splitted in selection medium in a 1:10 ratio as follows: medium was removed, petri dish was washed with 5-10 mL DPBS at RT; 1 mL 0.25% Trypsin-EDTA was added and during the 3-5 min incubation time, gentle mechanical force was additionally applied to facilitate the detachment of the cells; trypsinization was stopped by adding 10 mL of prewarmed (at 37 °C) selection medium; cells were then seeded accordingly (1:10) into new petri dishes. After growing enough cells, L-WRN-conditioned medium was produced by first splitting the cells in DMEM-Dil. in a 1:3 ratio. This time, cells were not transferred into petri dishes but into T-175 flasks (660160, Greiner CELLSTAR®) containing 30 mL cell-DMEM-Dil.-solution. Flasks were incubated at 37 °C for 2-3 days until cells became confluent. After collecting the first batch of medium, further 30 mL DMEM-Dil. were added, flasks were incubated and after 24 h, the medium was collected. The collection procedure was repeated up to 3-4 times. After every collection, the medium was centrifuged at 18 °C and 2000 g for 5 min. Centrifuged media were pooled, filtered (Vacuum Filtration 1000 “rapid”-Filtermax with PES membrane 0.22 µm pore size, 99950, TPP) and stored at 4 °C. Wnt3A-conditioned medium was produced the same as WRN-medium with L-Wnt3A producing cells (ATCC® CRL2647™). Here, the selection medium was composed of DMEM-Dil and 0.4 mg/mL G418.

Ready-to-use growth medium was prepared by combining WRN (75%)- and Wnt3A (25%)-media and adding the following growth factors: 1x B27, 1x N2, 50 ng/mL EGF, 1 mM N-Acetylcystein (A7250, Sigma-Aldrich) previously dissolved in DPBS, and 10 µM Rock Inhibitor (Y-27632 dihydrochloride; 1254, biotechne®).

### Injection needles for microinjection

Injection needles (ca. 5 cm) were made from glass capillaries (1.0 mm OD x 0.78 mm ID, W3 30-0036, Harvard Apparatus) pulled on a Flaming/Brown P97 Micropipette puller (Sutter Instrument, Novato CA 94949) under the following settings: Heat: 761; Pull: 80; Vel: 70; Del/Time: 170; Pressure: 493. The tip of the freshly made needles was opened by gently pinching the needles through a tissue previously soaked in 70 % EtOH. Injection needles were prepared at least one day prior to microinjection.

To estimate the volume of liquid per injection shot (V_s_), 3 µL trypan blue were loaded into a microinjection needle using a microloader tip (5242956003, Eppendorf). The needle was inserted into the pen of the microinjector and the content of the needle was released into a petri dish filled with DPBS until the needle was empty. V_s_ was calculated by dividing the volume loaded in the needly by the number of shots needed to empty it. The estimation of V_s_ was performed three times and the average V_s_ was 39.7 nL.

### Preparation for microinjection of organoids

In the passage previous to microinjection, organoid fragments were seeded in a 4-well plate (144444, Thermo Fisher Scientific). If histology of the organoids was planned, the wells of the 4-well plate additionally contained a glass slide (83.1840.002, SARSTEDT).

Organoids were microinjected with colibactin-producing *E. coli* (CoPEC) and colibactin-non-producing *E. coli* (noCoPEC). *E. coli* strain M1/5 *rpsL*K42R-*gfp* (CoPEC) is a variant of the WT *E. coli* B2 pks+ M/15 human fecal isolate. In the isogenic mutant strain M1/5 *rpsL*K42R-*gfp* Δ*clbR* (noCoPEC), the *clbR* gene in the pks island was deleted, hindering the bacterium’s ability to produce colibactin. Both *E. coli* strains carry the *rpsL*K42R mutation, which confers them resistance to streptomycin^83^. The *gfp* construct was additionally incorporated into the bacterial chromosome at the chromosomal attachment site of bacteriophage lambda (λ-*attB*) via recombination as described by Datsenko and Wanner^97^. *E. coli* strains were grown in Lysogeny Broth (LB) and aliquots were stored at −80 °C in LB 10% glycerol until usage. The following viable cell count of the glycerol stocks was determined by plating: CoPEC (4.1×10^8^ CFU/mL) and noCoPEC (3.7×10^8^ CFU/mL).

On the day of the microinjection, the glycerol stocks (500 µL) were thawed at RT. After centrifugation at 350 g and 4 °C for 5 min, the supernatant was discarded, and the bacterial pellet was resuspended in 500 µL Advanced DMEM/F12 medium at 4.1×10^8^ CFU/mL for CoPEC and 3.7×10^8^ CFU/mL for noCoPEC, and stored on ice. The bacteria solution for the microinjection was prepared under sterile conditions by mixing blue liquid food colorant (Backund Speisefarbe Blau, Rosenheimer Gourmet Manufaktur) with the bacterial solution in a 1:10 ratio.

### Microinjection of bacteria into the organoids

The construct for the microinjection, consisting of a microscope, mechanical arm, and injection system was installed under the sterile bench. Injection needles and further labware were also placed under the sterile bench and irradiated with UV light for 15 min.

The injection needle was loaded with 7 µL bacteria solution using a microloader tip and inserted into the injection pen of the microinjector (FemtoJet® 4i, Eppendorf). The injection pen was subsequently fixed to the mechanical arm attached to the microscope. Before using the needle for organoid-injection, the needle was tested for liquid release by injecting the stained bacteria solution into a small petri dish filled with Advanced DMEM/F12 medium. Liquid was released from the needle when pressing the foot pedal of the microinjector adjusted to the following settings: 30-80 hPA pressure (Injection); 0.3-0.6 sec (Time); 0 hPA pressure (Compensation). If the needle did not clog, its content sufficed for approximately 2-3 wells. Considering this, the volume of bacteria solution injected into the organoids of each well was approximately 3 µL.

For the microinjection of the organoids, the 4-well plate containing the organoids was positioned on the microscopés sample table. The position of the injection pen was adjusted using the mechanical arm, the needle was inserted into the organoids and the bacterial content was released by pressing the foot pedal of the microinjector. After injecting as many organoids per well as possible, an image of every well was taken with the Zeiss Stemi 508 stereomicroscope (Zeiss), and the growth medium was exchanged for Gentamicin-containing growth medium (10 µg/mL). The microinjection of noCoPEC, CoPEC and FC (in that order) was performed the same day. 2-3 4-well plates of organoids were microinjected with each condition.

### Assessment of phenotype via histology

Glass slides were placed in the 4-well plates, where organoids were seeded in MG the passage prior to fixation. The day of fixation, growth medium was removed and replaced by prewarmed (37°C) 500 µL PBS+, which were afterwards removed and replaced by prewarmed (37°C) 500 µL 4% PFA. After a 30-min incubation, 4% PFA was carefully discarded and replaced by 250 µL PBS+. The glass slides were carefully removed from the wells using tweezers and a needle. Each glass slide was placed into a histology cassette. After fixation, organoids were subsequently subjected to overnight dehydration in increasing concentrations of EtOH, xylene, and paraffin in a tissue processing unit (Leica TP1020). The day after, organoids were carefully embedded in paraffin by combining 4 wells into one paraffin block. When needed, Formalin-Fixed Paraffin-Embedded (FFPE) blocks containing organoids were precooled at approximately −15 °C on a cooling plate (CP-4D, KUNZ Instruments), cut into 2.5 µm sections with the microtome (RM2245, Leica), transferred to a 45 °C warm water bath to straighten until mounting on microscope slides (631-0108, VWR® SuperFrost®). Sections were air-dried overnight at room temperature and stored in light-protecting boxes until downstream processing. To increase tissue attachment of the sections to the glass, slides were heated at 60 °C for 60 min prior to staining. Stained slides were scanned and evaluated in the freely available software QuPath (Version 0.2.3.). For the Ki67- and the γH2AX-evaluation, the positive cell detection option was used with a manually adjusted, slide-dependent threshold. Organoids of the slide were first defined as objects. Positive and total cell nuclei were automatically detected in the previously defined objects.

### Immunohistochemistry (IHC)

IHC was performed in glass containers and in a moist, light-protected chamber. For incubation steps, tissue on the slides was completely overlaid with 200 µl of the respective solutions. Washing buffer TBST (TBS; 1244.1, Roth) with 0.05 % TWEEN-20 (P2287, Sigma-Aldrich), 3 % H_2_O_2_ (1.07209.0250, Merck) in DPBS and 3 % BSA (0163.2, Roth) in TBST were prepared before staining.

FFPE sections were deparaffinized in Roti-Histol (2× 10 min; 6640.1, Roth) and dehydrated with graded alcohol (2× 100 % for 5 min, 2x 96 % for 2 min, 2x 70 % for 2 min; Roth). After antigen retrieval with citrate buffer (pH=6, 10x, C9999, Sigma-Aldrich) in a pressure cooker (2100-Retriever, aptum), a hydrophobic barrier was drawn around the tissue with a PAP-pen (VEC-H-4000, Vector Laboratories). Sections were then washed with TBST (3x 5 min), and the endogenous peroxidase activity was quenched using 3 % H_2_O_2_ /PBS (10 min at RT). After another washing step (3x 5 min with TBST), the sections were blocked with 3 % BSA/TBST + 5 % goat serum (S-1000-20, Vector Laboratories) + streptavidin (1 h at RT; SP-2002, Vector Laboratories). Following incubation with the 1^st^ AB (Table 1) in 3 % BSA/TBST + biotin (1-2 h at RT; SP-2002, Vector Laboratories) and subsequent washing, the sections were incubated with the 2^nd^ AB (Goat Anti-Rabbit IgG Antibody (H+L), Biotinylated, BA-1000-1.5, Vector Laboratories) in 3 % BSA/TBST for 30 min at RT and washed again. For increasing the signal intensity, sections were incubated with the avidin-biotin-complex (VECTASTAIN® Elite® ABC-HRP Kit, PK-6100, Vector Laboratories) for 30 min at RT and were subsequently washed. For signal detection, the sections were incubated with 3,3’-diaminobenzidine (DAB Substrate Kit, SK-4100) for 2.5 min and washed briefly with distilled water. Cell nuclei were labeled with haematoxylin (Gil III, 25099, Waldeck) for 20-40 s. Sections were the rehydrated with graded alcohol (2x 70 % for 2 min, 2x 96 % for 2 min, and 2× 100 % for 5 min, Roth) and immersed Roti-Histol (2× 10 min). After completion, stained slides were mounted in Pertex embedding medium (00811-EX, HistoLab) and covered with coverslips (11911998, MENZEL-Gläser).

**Table 1:**
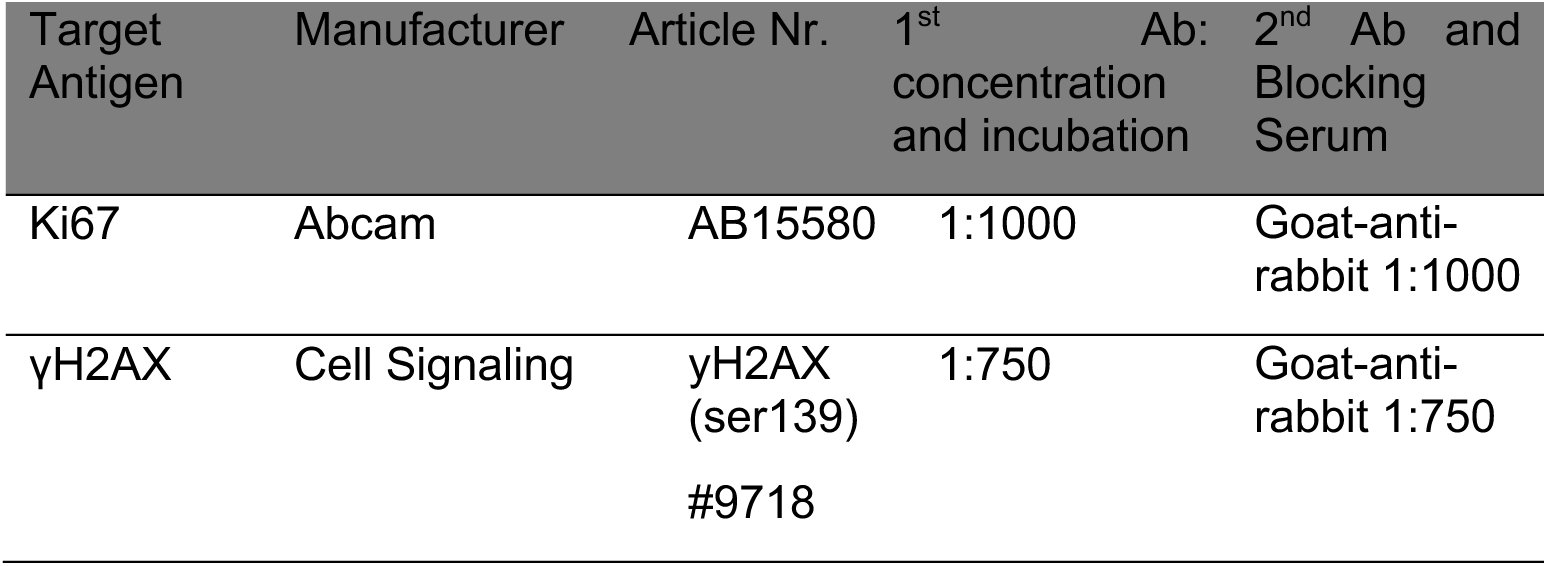
Specifications of primary antibodies for Immunohistochemistry.

### Plate preparation and single-cell sorting of organoids

The protocol of plate-based scRNA-seq was based on the protocol of Sagar et al. ^98^. Pipetting steps during plate preparation were performed with the I.DOT (DISPENDIX, Stuttgart). The wells of a 384-well PCR plate (AB-1384, Thermo Fisher Scientific) were pre-loaded with 1 µL mineral oil (M8410, Sigma-Aldrich) per well. Lysis buffer was prepared considering the following mixture per well: 280 nL of 0.35 % Triton (T8787, Sigma-Aldrich), 80 nL of nuclease-free water (AM9937, Thermo Fisher Scientific) and 40 nL dNTPs (N0447S, NEB). In each well, lysis buffer was added together with 80 nL of a unique primer (1 µM)^98^. Each 384-well PCR plate consisted of two sets of 192 different unique primers from Integrated DNA Technologies (IDT, Leuven, Belgium). Primer sequence and method of primer dilution were described by Sagar et al. ^98^. Plates were then covered with an adhesive, aluminium foil (3911257, VWR), centrifuged at 4 °C and full speed for 1 min, and were stored at −20 °C until further use. Prior to cell sorting, plates were centrifuged at 4 °C and 2000 g for 5 min and placed on ice.

For cell sorting, organoids were digested into single cells and stained for viability. DPBS, 0.25% Trypsin-EDTA, and DMEM-Dil-solution were prewarmed at 37 °C. 350 µL of warm DPBS were added to each well of the 4-Well plate containing organoids. The content of the wells was mixed and transferred into a 15 mL collection tube. Organoids were fragmented by pipetting up and down (5-10x) the content of the tube with a self-made narrow-end glass pipette. After centrifuging the tubes at room temperature (RT) and 350 g for 3 min, the supernatant was discarded, and pellets were resuspended in 400-600 µL warm 0.25 % Trypsin-EDTA. During incubation at 37 °C for 21 min, the content of the tubes was mixed thoroughly 3 times. To stop trypsinization, DMEM-Dil was added to the tube until reaching 10 mL. At this stage, the tube was placed in the bead bath at 37 °C until further use. FACS buffer was prepared in advance as follows: DPBS + 2 mM EDTA (AM9260G, Invitrogen by Thermo Fisher Scientific) and 2 %/w BSA (0163.4, Roth). Staining solution composed of zombie NIR (BioLegend, San Diego, USA) and FACS buffer was freshly prepared in a 1:800 ratio. Tubes containing the digested organoids were centrifuged at RT and 350 g for 5 min. Supernatant was removed, pellets were resuspended in 1 mL DPBS, and transferred into 1.5 mL LoBind-tube (022431021, Eppendorf). After centrifugation and removal of the supernatant, pellets were resuspended in staining solution (300 µL staining solution for cells from 4 wells) and incubated for 20 min. After centrifugation and removal of the supernatant, pellets were resuspended in 300-500 µL FACS buffer and transferred into a new 1.5 mL LoBind tube.

Single, live cells of the organoids were sorted with the CytoFLEX SRT Cell Sorter (Beckman Coulter, Brea, USA) into the 384-well PCR plates previously prepared. After sorting, plates were covered with adhesive aluminum foil, centrifuged at 4 °C and 2000 g for 5 min, and stored at −80 °C until further use.

### Preparation for plate-based single-cell RNA sequencing (scRNA-seq)

384-well PCR plates containing single cells were then processed as follows:

#### Reverse transcription

384-well PCR plates containing the sorted cells were taken from the −80 °C freezer and centrifuged at 4 °C and full speed for 1 min. The plates were then placed in a thermal cycler (T100, Biorad/ Mastercycler nexus gradient, Eppendorf) for cell lysis at 72 °C for 5 min. After centrifugation (4 °C, 2000 rpm, 1 min), the plates were placed on ice until further use. For the 1^st^ strand synthesis, a master mix was prepared considering the following mixture per well: 160 nL Protoscript II reaction buffer (M0368L, NEB), 80 nL 0.1 M DTT (M0368L, NEB), 40 nL RNase inhibitor (M0314S, NEB), 40 nL Protoscript II reverse transcriptase (M0368L, NEB). The master mix was distributed into the wells of the plates with the I-DOT (DISPENDIX, Stuttgart). After centrifugation (4 °C, 2000 rpm, 1 min), plates were placed in the thermocycler for 1^st^ strand synthesis (42 °C for 1 h, 70 °C for 10 min, and 4 °C for unlimited time). For the 2^nd^ strand synthesis, a master mix was prepared considering the following mixture per well: 3.262 µL nuclease free water (AM9937, Thermo Fisher Scientific), 0.462 µL second strand buffer (B6117S, NEB), 0.092 µL dNTPs (N0447S, NEB), 0.032 µL *E. coli* ligase (M0205S, NEB), 0.12 µL Rnase H (M0297S, NEB) and 0.032 µL DNA polymerase I (M0209S, NEB). Before and after adding the master mix with the I-DOT, plates were centrifuged at 4 °C and 2000 rpm for 1 min. Plates were placed again in the thermocycler for 2^nd^ strand synthesis (16 °C for 2 h and 4 °C for endless time). Of note, before every centrifugation step, plates were covered with adhesive aluminum foil.

#### Clean-Up of complementary DNA (cDNA)

AMPure beads (A63880, Beckman Coulter) were prewarmed to RT for at least 30 min and 80 % EtOH was freshly prepared with EtOH absolute (16685992, 200 Proof, Fisher BioReagents^TM^). After centrifuging the plate at 4 °C and 2000 rpm for 1 min, wells of the columns 1-6, 7-12, 13-18, 19-24 were pooled and transferred into 6×1.5 mL LoBind tubes. The content of each LoBind tube was distributed in 4 wells of a 96-well PCR LoBind plate. AMPure beads were vortexed until well dispersed, added (0.8x), and mixed well with the samples on the plate. After incubation for 10 min at RT, the plate was placed on a magnet for 2 min until the solution became clear. Clear supernatant was removed and discarded. Remaining pellets were washed twice by adding 360 µL 80 % EtOH for 30 s. After removing the 80 % EtOH from the 2^nd^ washing step, pellets were left to dry for 2-3 min on the magnet. The plate was removed from the magnet and the pellets of each pool were resuspended together with 14 µL in nuclease-free water. After a short incubation of 2 min at RT, the plate was placed again on the magnet until the solution became clear. For proceeding with *in-vitro* transcription, 12.8 µL of the clear solution from each pool were transferred to the tube of a PCR tube strip. At this stage, samples were stored at −20 °C until further processing.

#### In-vitro transcription (IVT)

A master mix (MM) was prepared with the reagents of the MegaScript T7 kit (AM1333, Invitrogen) considering the following mixture per sample (tube of the PCR tube strip): 3.2 µL ATP, 3.2 µL GTP, 3.2 µL CTP, 3.2 µL UTP, 3.2 µL T7 buffer (10x), 3.2 µL T7 enzyme. 19.2 µL IVT MM were added and mixed well with each sample. After a short spin-down of the tube strip, the IVT program of the thermocycler was initiated: 37 °C for 13 h and 4 °C for an endless time.

#### Clean-up of amplified RNA (aRNA)

RNAclean XP beads (A63987, Beckman Coulter) were prewarmed to RT for at least 30 min and 70 % EtOH was freshly prepared with ethanol absolute (16685992, 200 Proof, Fisher BioReagents^TM^). For the initial cleaning step, 12 µL of EXO-SAP (78200.200.UL, Thermo Fisher Scientific) were added and mixed well with each sample, and the following program was initiated in the thermocycler: 37 °C for 15 min and 4 °C for an endless time. For the fragmentation step, 4.88 µL of 10x fragmentation buffer (E6150S, NEB) were added and mixed well with each sample, and the following program was initiated in the thermocycler: 94 °C for 3 min and 4 °C for an endless time. Samples were placed on ice and 4.88 µL of 10x fragmentation stop buffer (E6150S, NEB) were added and mixed well with each sample. RNAclean XP beads were vortexed until well dispersed, added (1.2x) and mixed well with each sample. After incubation for 10 min at RT, the PCR tube strip containing the samples was placed on a magnet until the solutions became clear. Clear supernatants were removed and discarded. Remaining pellets were washed (3x) by adding 360 µL 70 % EtOH for 30 s. After removing the 70 % EtOH from the 3^rd^ washing step, pellets were left to dry for 2-3 min on the magnet. Pellets were then resuspended in 7 µL in nuclease-free water (AM9937, Thermo Fisher Scientific). After a short incubation of 2 min at room temperature, the plate was placed again on the magnet until the solution became clear. 6.5 µL of every sample were transferred into a new tube and placed on ice until further use. RNA concentration of the samples was measured with the Bioanalyzer (2100, Agilent) using the Agilent RNA 6000 Pico kit (5067-1513, Agilent).

#### Library preparation

1 µL random HexRT primer (10 µM, IDT, Supplementary table) was mixed with 0.5 µL dNTPs and added to 5 µL aRNA. After incubation at 65 °C for 5 min, samples were placed on ice. 1^st^ strand MM was prepared considering the following mixture per sample: 2 µL Protoscript II reaction buffer (M0368L, NEB), 1 µL DTT (0.1 M, M0368L, NEB), 0.5 µL RNase inhibitor (M0314S, NEB), 0.5 µL Protoscript II reverse transcriptase (M0368L, NEB). 1^st^ strand MM was added to every sample and the following program was initiated in the thermocycler: 25 °C for 10 min and 42 °C for 1h.

#### PCR amplification

PCR MM was prepared with the following mixture per sample: 11 µL water, 25 µL HF PCR MM (M0541S, NEB), 2 µL Illumina primer RP1 (fw, 10 µM, IDT, Supplementary table). In addition to the 38 µL PCR MM, 2 µL indexed RPIx (i7) primer (IDT, Supplementary table) were added to each sample. PCR was performed in the thermocycler as follows: preheating to 98°C; initial denaturation (98 °C, 30 s); 11-15 cycles of denaturation (98 °C, 10 s); annealing (60 °C, 30 s); elongation (72 °C, 30 s); final elongation (72 °C, 10 min) and cooling to 4 °C for endless time.

#### PCR clean-up

AMPure beads were prewarmed to RT for at least 30 min, and 80 % EtOH was freshly prepared. AMPure beads were vortexed until well dispersed, added (0.9x), and mixed well with each PCR reaction. After incubation for 10 min at RT, the PCR tube strip was placed on a magnet until the solutions became clear. Clear supernatant was removed and discarded. Remaining pellets were washed twice by adding 180 µL 80 % EtOH for 30 s. After removing the 80 % EtOH from the 2^nd^ washing step, pellets were left to dry for 2-3 min on the magnet. Pellets were resuspended in 26 µL nuclease-free water (AM9937, Thermo Fisher Scientific). After a short incubation of 2 min at RT, the PCR tube strip was placed again on the magnet until the solution became clear. 25 µL of the clear solutions were transferred into a new DNA LoBind tube. Quality control of the libraries was performed with the Qubit (Invitrogen by Thermo Fisher Scientific) and Bioanalyzer (5067-4626, Agilent). Finished libraries were stored at −20 °C until shipment to Biomarker Technologies GmbH (BMK, Münster, Germany) for scRNA-seq.

### Sequencing of scRNA-seq libraries and data analysis

Single-cell RNA sequencing was performed using a miniaturized version of CEL-Seq2 protocol^99^. 48 libraries with 96 cells each were prepared separately, pooled together and sequenced on one lane of Illumina Nova PE150 sequencer at a depth of minimum 100,000 reads/cell. Indexing of reference genome and quantification of gene expression counts were performed using simpleaf^100^ (v0.17.2) and alevin-fry^101^ (v0.10.0). First, a spliced + intronic reference was created using the simpleaf index command, the genome assembly GRCm39 by the Genome Reference Consortium, and setting the read length to 150 base pairs. Next, the quantification was performed using the simpleaf quant command with the UMI resolution mode “parsimony”. As this was a custom library preparation protocol with known barcodes, a custom chemistry “{u[6]b[6]x:}2{r:}” and a list of possible barcodes were supplied.

Data analysis was performed using Python (v3.12.4) and scanpy (v1.10.2). All 48 libraries were combined, and cells with fewer than 300 genes or 500 UMI counts were excluded as low-quality cells. Importantly, ribosomal genes (small and large subunits), mitochondrial genes and genes with *Gm*-identifier were excluded from the analysis. Normalization was performed using the normalize_total function, followed by log1p-transformation and scaling. PCA and UMAP were calculated using default settings. Clustering was performed with the Leiden algorithm with a resolution of 0.4 while setting the flavor to “igraph” and n_iterations to 2. The cell cycle status was determined using the score_genes_cell_cycle function and the gene sets published by Kowalczyk *et al.*^102^. GSEA was performed using decoupler (v1.7.1).

### Statistics

Unless stated otherwise, the statistical analysis was performed using GraphPad Prism 10.2.2. for Windows. Comparison of more than two groups per data set was performed first by testing each data set for normality using the Shapiro-Wilk method and assessing the difference between the variance (SD) of the groups using the Brown-Forsythe test. If data passed the normality test and the SDs were assumed to be equal, an Ordinary one-way ANOVA with Tukeýs multiple comparisons test was performed. If the data passed the normality test and the SDs were significantly different, a Brown-Forsythe and Welch ANOVA with Dunnett’s T3 multiple comparisons test was performed. If the data did not pass the normality test, a Kruskal-Wallis with Dunńs multiple comparisons test was performed. Plots are depicted as Mean with SEM.

### Use of Artificial Intelligence (AI)

AI tools were used for a first screening of the genes and pathways found to be over-/underexpressed or up-/downregulated in the scRNA-seq data. After gaining a first overview, the function of the genes and pathways was verified in the literature before describing the results and drawing any conclusions. The AI tools used for this purpose were Perplexity and Elicit. In addition, ChatGPT and the AI-Assistant of DeepL were used for correcting grammar and optimizing word choice.

## Supporting information

Sequences of primers used for PCR amplification when preparing the organoid cells for single-cell RNA sequencing

## Authors contributions

**Study, concept and design:** Michael Quante, Julian A. Abrams

**Acquisition of data:** Andrea Proaño-Vasco, Lauren Houle, Yiwei Sun

**Analysis and interpretation of data:** Andrea Proaño-Vasco, David Obwegs, Sagar, Lauren Houle, Ceylan Tanes

**Drafting the manuscript:** Andrea Proaño-Vasco, Michael Quante, Julian A. Abrams

**Critical revision of the manuscript for important intellectual content:** all authors

**Statistical analysis:** Andrea Proaño-Vasco, Lauren Houle

**Obtained funding:** Michael Quante, Julian A. Abrams, Sagar,

**Technical or material support:** Bärbel Stecher, Ulrich Dobrindt, Kerstin Bruder, Céline Ritzkowski, Sophia Eichner, Lioba Klaas, Linus R. Schömig, Donja Sina Mohammed-Shahi,

**Study supervision:** Michael Quante, Julian A. Abrams

## Data availability

The primary read files and the raw counts as well as processed data for all single-cell RNA sequencing datasets reported in this paper are available upon request to corresponding author. All other raw data are available upon request to corresponding author-

## Code availability

Codes to reproduce the data analysis and figures are available upon request to corresponding author.

