## Supplementary material for "Colibactin-producing *E. coli* promote carcinogenesis of gastroesophageal adenocarcinoma and simultaneously induce autophagy and differentiation": Sequences of primers used for PCR amplification when preparing the organoid cells for single-cell RNA sequencing

***Supplementary Table 1****: Sequences of primers used for PCR amplification when preparing the organoid cells for single-cell RNA sequencing*

| | |
|---|---|
| random HexRT | GCCTTGGCACCCGAGAATTC- CANNNNNN |
| RP1 | AATGATACGGCGACCACCGAGATCTACACGTTCAGAGTTC TACAGTCCGA |
| RPI1 | CAAGCAGAAGACGGCATACGAGAT***CGTGAT***GTGACTGGAGTTCCTTGGCACCCGAGAATTCCA |
| RPI2 | CAAGCAGAAGACGGCATACGAGAT***ACATCG***GTGACTGGAGTTCCTTGGCACCCGAGAATTCCA |
| RPI3 | CAAGCAGAAGACGGCATACGAGAT***GCCTAA***GTGACTGGAGTTCCTTGGCACCCGAGAATTCCA |
| RPI4 | CAAGCAGAAGACGGCATACGAGAT***TGGTCA***GTGACTGGAGTTCCTTGGCACCCGAGAATTCCA |
| RPI5 | CAAGCAGAAGACGGCATACGAGAT***CACTGT***GTGACTGGAGTTCCTTGGCACCCGAGAATTCCA |
| RPI6 | CAAGCAGAAGACGGCATACGAGAT***ATTGGC***GTGACTGGAGTTCCTTGGCACCCGAGAATTCCA |
| RPI7 | CAAGCAGAAGACGGCATACGAGAT***GATCTG***GTGACTGGAGTTCCTTGGCACCCGAGAATTCCA |
| RPI8 | CAAGCAGAAGACGGCATACGAGAT***TCAAGT***GTGACTGGAGTTCCTTGGCACCCGAGAATTCCA |
| RPI9 | CAAGCAGAAGACGGCATACGAGAT***CTGATC***GTGACTGGAGTTCCTTGGCACCCGAGAATTCCA |
| RPI10 | CAAGCAGAAGACGGCATACGAGAT***AAGCTA***GTGACTGGAGTTCCTTGGCACCCGAGAATTCCA |
| RPI11 | CAAGCAGAAGACGGCATACGAGAT***GTAGCC***GTGACTGGAGTTCCTTGGCACCCGAGAATTCCA |
| RPI12 | CAAGCAGAAGACGGCATACGAGAT***TACAAG***GTGACTGGAGTTCCTTGGCACCCGAGAATTCCA |
| RPI13 | CAAGCAGAAGACGGCATACGAGAT***TTGACT***GTGACTGGAGTTCCTTGGCACCCGAGAATTCCA |
| RPI14 | CAAGCAGAAGACGGCATACGAGAT***GGAACT***GTGACTGGAGTTCCTTGGCACCCGAGAATTCCA |
| RPI15 | CAAGCAGAAGACGGCATACGAGAT***TGACAT***GTGACTGGAGTTCCTTGGCACCCGAGAATTCCA |
| RPI16 | CAAGCAGAAGACGGCATACGAGAT***GGACGG***GTGACTGGAGTTCCTTGGCACCCGAGAATTCCA |
| RPI17 | CAAGCAGAAGACGGCATACGAGAT***CTCTAC***GTGACTGGAGTTCCTTGGCACCCGAGAATTCCA |
| RPI18 | CAAGCAGAAGACGGCATACGAGAT***GCGGAC***GTGACTGGAGTTCCTTGGCACCCGAGAATTCCA |
| RPI19 | CAAGCAGAAGACGGCATACGAGAT***TTTCAC***GTGACTGGAGTTCCTTGGCACCCGAGAATTCCA |
| RPI20 | CAAGCAGAAGACGGCATACGAGAT***GGCCAC***GTGACTGGAGTTCCTTGGCACCCGAGAATTCCA |
| RPI21 | CAAGCAGAAGACGGCATACGAGAT***CGAAAC***GTGACTGGAGTTCCTTGGCACCCGAGAATTCCA |
| RPI22 | CAAGCAGAAGACGGCATACGAGAT***CGTACG***GTGACTGGAGTTCCTTGGCACCCGAGAATTCCA |
| RPI23 | CAAGCAGAAGACGGCATACGAGAT***CCACTC***GTGACTGGAGTTCCTTGGCACCCGAGAATTCCA |
| RPI24 | CAAGCAGAAGACGGCATACGAGAT***GCTACC***GTGACTGGAGTTCCTTGGCACCCGAGAATTCCA |
| RPI25 | CAAGCAGAAGACGGCATACGAGAT***ATCAGT***GTGACTGGAGTTCCTTGGCACCCGAGAATTCCA |
| RPI26 | CAAGCAGAAGACGGCATACGAGAT***GCTCAT***GTGACTGGAGTTCCTTGGCACCCGAGAATTCCA |
| RPI27 | CAAGCAGAAGACGGCATACGAGAT***AGGAAT***GTGACTGGAGTTCCTTGGCACCCGAGAATTCCA |
| RPI28 | CAAGCAGAAGACGGCATACGAGAT***CTTTTG***GTGACTGGAGTTCCTTGGCACCCGAGAATTCCA |
| RPI29 | CAAGCAGAAGACGGCATACGAGAT***TAGTTG***GTGACTGGAGTTCCTTGGCACCCGAGAATTCCA |
| RPI30 | CAAGCAGAAGACGGCATACGAGAT***CCGGTG***GTGACTGGAGTTCCTTGGCACCCGAGAATTCCA |
| RPI31 | CAAGCAGAAGACGGCATACGAGAT***ATCGTG***GTGACTGGAGTTCCTTGGCACCCGAGAATTCCA |
| RPI32 | CAAGCAGAAGACGGCATACGAGAT***TGAGTG***GTGACTGGAGTTCCTTGGCACCCGAGAATTCCA |
| RPI33 | CAAGCAGAAGACGGCATACGAGAT***CGCCTG***GTGACTGGAGTTCCTTGGCACCCGAGAATTCCA |
| RPI34 | CAAGCAGAAGACGGCATACGAGAT***GCCATG***GTGACTGGAGTTCCTTGGCACCCGAGAATTCCA |

| | |
|---|---|
| RPI35 | CAAGCAGAAGACGGCATACGAGAT***AAAATG***GTGACTGGAGTTCCTTGGCACCCGAGAATTCCA |
| RPI36 | CAAGCAGAAGACGGCATACGAGAT***TGTTGG***GTGACTGGAGTTCCTTGGCACCCGAGAATTCCA |
| RPI37 | CAAGCAGAAGACGGCATACGAGAT***ATTCCG***GTGACTGGAGTTCCTTGGCACCCGAGAATTCCA |
| RPI38 | CAAGCAGAAGACGGCATACGAGAT***AGCTAG***GTGACTGGAGTTCCTTGGCACCCGAGAATTCCA |
| RPI39 | CAAGCAGAAGACGGCATACGAGAT***GTATAG***GTGACTGGAGTTCCTTGGCACCCGAGAATTCCA |
| RPI40 | CAAGCAGAAGACGGCATACGAGAT***TCTGAG***GTGACTGGAGTTCCTTGGCACCCGAGAATTCCA |
| RPI41 | CAAGCAGAAGACGGCATACGAGAT***GTCGTC***GTGACTGGAGTTCCTTGGCACCCGAGAATTCCA |
| RPI42 | CAAGCAGAAGACGGCATACGAGAT***CGATTA***GTGACTGGAGTTCCTTGGCACCCGAGAATTCCA |
| RPI43 | CAAGCAGAAGACGGCATACGAGAT***GCTGTA***GTGACTGGAGTTCCTTGGCACCCGAGAATTCCA |
| RPI44 | CAAGCAGAAGACGGCATACGAGAT***ATTATA***GTGACTGGAGTTCCTTGGCACCCGAGAATTCCA |
| RPI45 | CAAGCAGAAGACGGCATACGAGAT***GAATGA***GTGACTGGAGTTCCTTGGCACCCGAGAATTCCA |
| RPI46 | CAAGCAGAAGACGGCATACGAGAT***TCGGGA***GTGACTGGAGTTCCTTGGCACCCGAGAATTCCA |
| RPI47 | CAAGCAGAAGACGGCATACGAGAT***CTTCGA***GTGACTGGAGTTCCTTGGCACCCGAGAATTCCA |
| RPI48 | CAAGCAGAAGACGGCATACGAGAT***TGCCGA***GTGACTGGAGTTCCTTGGCACCCGAGAATTCCA |
